# eSPred: Explainable scRNA-seq Prediction via Customized Foundation Models and Pathway-Aware Fine-tuning

**DOI:** 10.1101/2025.05.14.654052

**Authors:** Liwen Sun, Qianyu Yang, Jingxuan Zhang, Wenbo Guo, Lin Lin

## Abstract

Single-cell RNA sequencing (scRNA-seq) has been widely used for studying cellular heterogeneity, but its use for subject-level prediction and clinical applications is still limited. We introduce eSPred, a customized foundation model designed for predictive analysis of scRNA-seq. It integrates cell-type information through a grouping strategy during pre-training and leverages pathway information to guide network flow during fine-tuning. Across multiple datasets, eSPred improves prediction accuracy and highlights pathways linked to disease mechanisms. These results suggest that eSPred can help bridge the gap between single-cell data and subject-level clinical insights, supporting more precise diagnosis and better-informed treatment decisions.

## 1 Introduction

Single-cell RNA sequencing (scRNA-seq) has revolutionized cellular and molecular biology by enabling high-resolution characterization of cell states and dynamics at an unprecedented scale [1]. Unlike bulk RNA sequencing, which averages gene expression across heterogeneous cell populations, scRNA-seq captures transcriptomic profiles at the individual cell level, allowing researchers to dissect cellular heterogeneity, identify rare cell types, and uncover lineage relationships [2–4]. This capability has advanced our understanding of complex biological processes, including development, disease progression, and immune responses, making scRNA-seq an indispensable tool in biomedical research [5–8].

Traditional scRNA-seq analysis methods have evolved to address the challenges of high dimensionality and sparsity inherent in single-cell data. Early approaches focused on preprocessing steps such as quality control, normalization, and batch effect correction [9, 10], which are crucial to ensuring data integrity. To manage the complexity of single-cell transcriptomic data, various dimensionality reduction techniques have been developed to enhance visualization and facilitate the identification of distinct cell populations [11–21]. In parallel, clustering algorithms classify cells based on transcriptomic profiles, enabling the discovery of biologically relevant subtypes [22–36]. To further dissect cellular states, differential expression analysis methods such as edgeR [37], EBSeq [38], and DESeq2 [39] identify genes associated with specific cell populations or conditions. Additionally, pathway analysis tools—including CellTICS [40], P-NET [41], SCENIC [42], PAGODA [43], GSVA [44], Enrichr [45], and MAST [46]—enhance biological interpretability by linking gene expression patterns to functional pathways. Together, these methodological advancements have significantly improved our ability to characterize cellular heterogeneity, infer regulatory mechanisms, and deepen our understanding of cellular functions and disease mechanisms.

Despite advancements in single-cell analysis, translating scRNA-seq findings into clinically relevant predictions remains a significant challenge. Accurate subject-level predictions are essential for clinical applications, yet existing methods struggle with the high dimensionality of single-cell data, limited sample sizes in clinical studies, and the need to extract meaningful subject-level patterns while preserving cellular heterogeneity. Existing methods for subject-level prediction fall into two main categories: regression-based approaches and deep learning techniques. Regression-based methods, such as Cell Population Mapping (CPM) [47], linear mixed-effects models (LMMs) [48], and pseudobulk aggregation [49] offer interpretability but often oversimplify the complexity of single-cell data. For instance, pseudobulk approaches aggregate gene expression across cells, effectively averaging out cell-type-specific signals and obscuring cellular heterogeneity [50]. Similarly, LMMs rely on unsupervised clustering to define cellular subgroups, introducing variability due to arbitrary cluster assignments and sensitivity to hyperparameters, such as resolution thresholds in graph-based clustering [51]. These limitations can lead to biased effect estimates and reduced predictive power in complex biological systems. Additionally, many of these methods apply dimensionality reduction techniques to alleviate computational burdens, which may inadvertently discard critical biological signals [52].

Deep learning techniques have emerged as a powerful alternative for subject-level prediction, demonstrating superior capacity to model nonlinear relationships in single-cell data. Methods such as DeepGeneX [53], ScRAT [54], ProtoCell4P [55], scGNN [56], scVAE [57], scPhere [58], scScope [59], and scDeepCluster [35] excel at capturing intricate dependencies within the data. However, their clinical applicability remains constrained by three key limitations: the potential loss of fine-grained cellular heterogeneity, sensitivity to small sample sizes, and a lack of biological interpretability. First, many deep learning models compromise cellular heterogeneity by relying on averaging or summarization techniques. For example, scVAE and scDeepCluster use Gaussian mixture models to cluster cells, implicitly assuming discrete cell states and overlooking transitional or mixed cell populations. Similarly, pseudobulk-inspired architectures like ProtoCell4P average cell-level features into subject-level summaries, masking rare subpopulations [50]. Second, deep learning methods often struggle with limited sample sizes and variable cell counts. Graph-based approaches such as scGNN require fixed-size cell graphs per sample, necessitating subsampling or imputation that distorts biological variation. Meanwhile, scalable models like scScope and scPhere demand extensive hyperparameter tuning—such as graph neighborhood size and latent dimensionality—to mitigate overfitting, a challenge exacerbated in small clinical datasets [60]. Finally, limited biological interpretability hinders the clinical adoption of deep learning models. While methods such as DeepGeneX and ScRAT generate predictions, they do not explicitly link learned features to biological pathways or regulatory networks. Although ScRAT integrates attention mechanisms to highlight influential genes, it fails to contextualize them within established functional pathways, reducing their clinical relevance.

Building on deep learning approaches, transformer-based foundation models have recently emerged as effective tools for scRNA-seq analysis. Models such as scBERT [61], Geneformer [62], scGPT [63], and scFoundation [64] leverage large-scale pretraining on diverse single-cell datasets. Using self-supervised learning and attention mechanisms, they capture gene interactions both within and across cells. Unlike traditional methods that treat genes as independent features, transformers analyze the entire transcriptome simultaneously, generating gene embeddings that account for transcriptional context and cell embeddings that preserve cellular heterogeneity [65]. They have demonstrated strong performance in cell-type annotation, batch effect correction, zero-shot transfer learning, and feature representation [61–63]. Their ability to identify subtle gene expression patterns and rare cell populations surpasses conventional clustering and dimensionality reduction methods. However, extending transformer-based foundation models to subject-level prediction presents challenges similar to those encountered with traditional deep learning methods—an area that remains largely unexplored.

To address this gap, we introduce eSPred, a framework that adapts foundation models for scRNA-seq analysis through three key innovations: (1) a cell-type-informed input strategy that preserves biological heterogeneity to support learning of discriminative patterns across diverse cell populations; (2) a pathway-guided cell embedding mechanism, adapted from CellTICS [40] and P-NET [41], which incorporates established biological pathways to enhance interpretability; and (3) a hierarchical prediction approach that links cell-level features to subject-level outcomes. Together, these components enable eSPred to overcome key limitations in subject-level prediction: it maintains cellular heterogeneity without oversimplified averaging, accommodates varying sample sizes without extensive hyperparameter tuning, and enhances biological insight through pathway-informed representations.

## 2 Results

### Overview of the eSPred model

As illustrated in Figure 1, the eSPred framework integrates three key components to bridge cellular heterogeneity with clinical outcome prediction: (1) a customized foundation model that employs a cell-type-informed input strategy during pre-training to generate *cell-type-aware gene embeddings*; (2) a pathway-integrated decoder that produces biologically interpretable *pathway-aware cell embeddings* during fine-tuning; and (3) a hierarchical classifier that links cell-level representations to subject-level clinical outcomes.

**Fig. 1:**
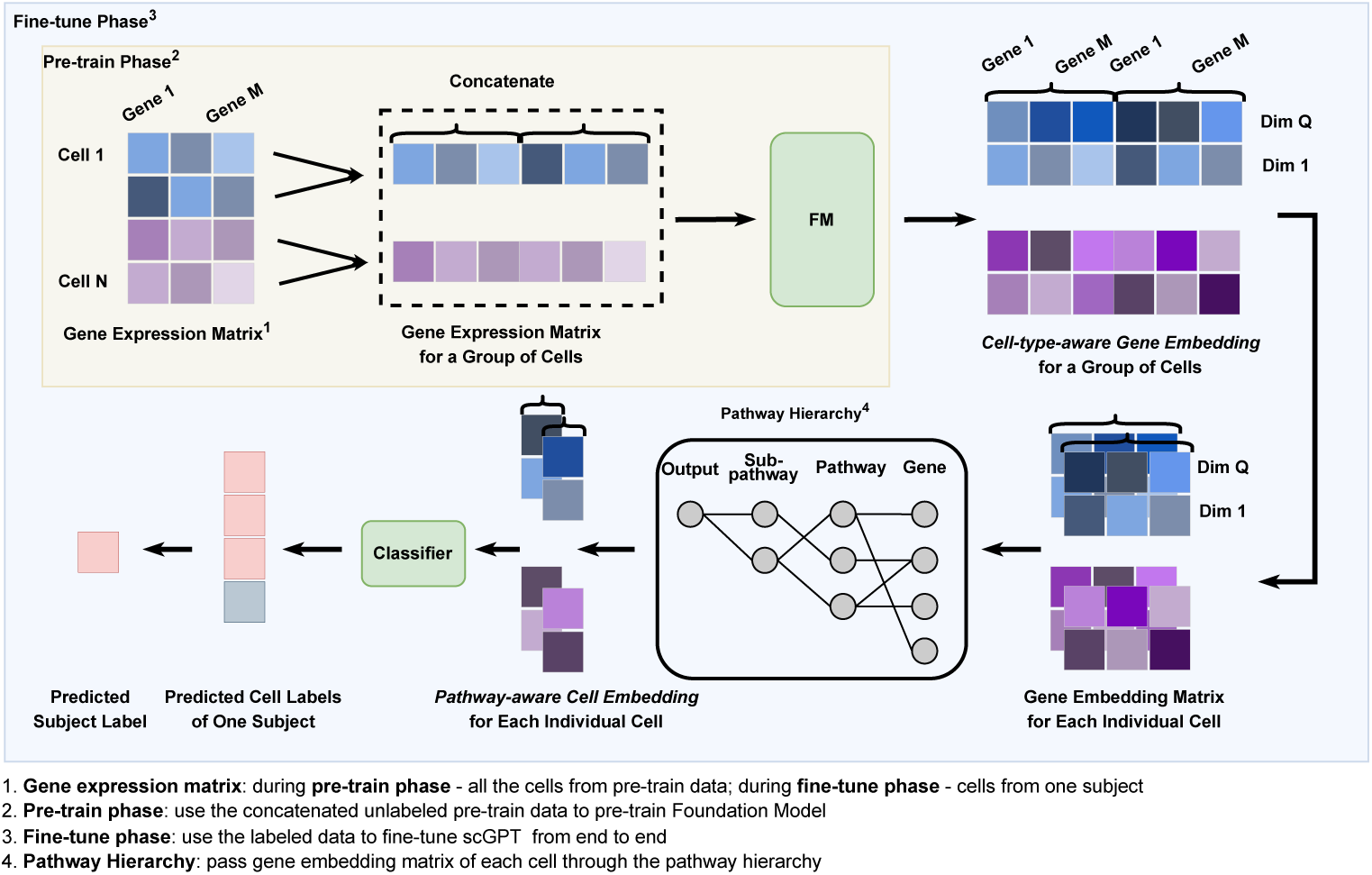
Overview of the proposed eSPred framework. The framework consists of three key stages: (1) Pre-training, where cells of the same type are organized into small groups, and their gene expression profiles are concatenated to create more informative training samples. These aggregated profiles are then fed into the foundation model to learn robust, high-level embeddings, which we name *Cell-type-aware Gene Embedding*. (2) Fine-tuning, where a pathway-aware mechanism integrates hierarchical pathway relationships from the Reactome database, ensuring that each gene connects only to its relevant pathways for biologically meaningful information propagation. We name the output as *Pathway-aware Cell Embedding*. (3) Prediction, where pathway-informed cell embeddings generate cell-level predictions, which are subsequently aggregated via majority voting to derive subject-level classifications.

### Customized foundation models leveraging cell type information

Among current foundation models, scGPT demonstrates superior performance and scalability. Its architecture effectively mitigates batch effects and generates both gene and cell embeddings with strong generalization capabilities across diverse datasets, making it well-suited for clinical applications [66]. Pre-trained on an extensive corpus of over 10 million cells from 65 publicly available CELLxGENE datasets, scGPT captures a broader spectrum of biological variation compared to alternative models such as Geneformer (trained on 29.9 million cells but with reduced specificity for single-cell data) and scBERT (limited to 1 million cells from Panglao) [61–63]. Given these substantive advantages, we adopt scGPT as the foundation model for our framework. However, our methodology can be adapted to other foundation models, as discussed in the Discussion section.

Building on the strengths of scGPT, we implement a structured input strategy during pre-training to enhance the model’s ability to capture biologically meaningful patterns. Specifically, we extract 1,200 highly variable genes (HVGs) from the pre-training dataset, using scGPT’s default variance-stabilizing transformation, which stabilizes gene expression variance across a wide dynamic range—facilitating the robust identification of HVGs. To preserve cellular heterogeneity while maintaining computational efficiency, we systematically group four cells of the same type, producing a 4 × 1200 gene expression matrix (where 4 denotes the number of cells and 1,200 the number of selected genes). This configuration nearly saturates scGPT’s sequence capacity (approximately 5,000 tokens), as each cell contributes 1,200 gene features, resulting in a total of 4,800 tokens per input. While reducing the number of HVGs (e.g., to 800) could allow grouping of more cells per input, this would risk omitting critical gene-level variation essential for downstream prediction. HVGs are known to retain the most informative and biologically relevant signals, and overly restrictive gene selection may degrade the quality of the learned cell embeddings. Retaining 1,200 HVGs also aligns with scGPT’s original pretraining parameters, thereby minimizing the need for extensive re-tuning. To form these groups within each sample, we follow a structured grouping strategy. When annotated cell-type information is available, we directly group four cells of the same type. Otherwise, we apply automated clustering methods, such as Leiden clustering on PCA-reduced data [26, 67], to assign cells into groups. To maintain a consistent group size, any remaining cells that do not form a complete group of four are excluded from the analysis. These parameters can be adjusted when implementing alternative foundation models.

Next, the 4 × 1200 matrix is first flattened into a 1 × 4800 vector through rowwise concatenation of the gene expression profiles, which is then used as input to the foundation model. During pre-training, the foundation model is optimized using a masked gene expression reconstruction objective, allowing it to simultaneously learn gene–gene interactions and cell-type-specific relationships. This customized input strategy facilitates the acquisition of richer biological representations, establishing a robust foundation for subsequent fine-tuning.

For the fine-tuning phase, we retain the same grouping strategy: four cells of the same type from the same sample are combined, and their gene expression profiles are concatenated into a single input vector. Passing this input through the pre-trained foundation model produces a cell-type-aware gene embedding matrix of dimensions 512 × 4800, where 512 is the embedding dimension (denoted as *Q* in Figure 1). We then partition this matrix into four separate submatrices, each corresponding to an individual cell and yielding a 512 × 1200 gene embedding matrix per cell.

### Pathway-integrated decoder during fine-tuning phase

Next, we derive the cell-level embedding by processing each cell’s gene embedding matrix. A common approach is element-wise mean pooling, which simply averages the gene embeddings across all genes. Alternatively, weighted pooling computes a weighted average, assigning greater influence to genes deemed more important. In contrast, the original scGPT implementation adopts a more sophisticated strategy: it introduces a special [CLS] token at the beginning of the gene sequence. This token is propagated through the transformer layers alongside gene tokens, allowing the model to learn an optimal aggregation function during training. The final embedding corresponding to the [CLS] token is then extracted as the cell’s representation. While foundation models excel at learning feature-rich representations, these aggregation methods often lack biological interpretability due to their black-box nature [61]. To overcome this limitation, we introduce a pathway-integrated decoder that transforms the foundation model’s outputs into biologically meaningful representations during fine-tuning.

Our pathway-integrated decoder implements a hierarchical neural network architecture guided by the Reactome database [68]. The network mirrors biological organization: the input layer consists of genes annotated in Reactome, intermediate layers represent pathways and subpathways, and the output layer aggregates higher-level biological signals. Crucially, the network’s connectivity is biologically grounded, with edges restricted to known gene–pathway relationships. This structure enhances interpretability and reduces overfitting by preventing spurious associations [69], distinguishing our model from conventional fully connected architectures where any gene can influence any pathway.

Each cell’s gene embedding matrix (512 × 1200) is processed through the pathway-integrated decoder, producing a vector pathway-aware cell embedding of length 512. To assess the contribution of individual pathways to disease classification, we compute pathway importance scores based on activation patterns. For each pathway (represented by a specific neuron), we collect its activation values across all cells, group them by outcome label (e.g., disease vs. normal), and compute the element-wise mean within each group. The L2 norm of the difference between these group-specific activation vectors is then used as the pathway’s importance score. This provides a direct and interpretable measure of which biological pathways are most influential in driving the model’s predictions.

### Hierarchical classifier for clinical outcome predictions

Following the generation of pathway-aware cell embeddings, we implement subject-level prediction using a hierarchical classification strategy. eSPred employs a systematic two-step approach to aggregate cell-level heterogeneity into coherent subject-level outcomes. First, each cell’s pathway-aware embedding is individually classified using a linear layer optimized with cross-entropy loss. Then, sample-level predictions are determined through a threshold-based majority voting mechanism: a sample is classified as positive if the proportion of cells predicted as positive exceeds a predefined threshold, typically set at 50%. This hierarchical design preserves cellular resolution while enabling robust subject-level inference. By first operating at the cell level, the model maintains the biological heterogeneity that would otherwise be lost in bulk-averaging approaches. At the same time, aggregating cell-level predictions mitigates the effects of technical noise and outlier cells, improving the stability and reliability of subject-level classifications. The voting threshold is tunable and can be adjusted depending on the disease context and the desired sensitivity–specificity trade-off.

### eSPred enhances subject-level prediction performance

We evaluate eSPred’s predictive performance against both traditional machine learning models and deep learning methods developed for scRNA-seq data analysis. Specifically, we compare eSPred with ProtoCell4P and DeepGeneX, two specialized models for prediction and interpretation, alongside conventional classifiers including Logistic Regression (LR), Random Forest (RF), Support Vector Machine (SVM), and Multi-Layer Perceptron (MLP). Benchmarking is conducted across three disease datasets—COVID-19, Lupus, and Lung—with evaluations based on class-specific precision and recall for distinguishing disease from normal states. Details of the three datasets are provided in the **Dataset** section.

As shown in Figure 2, eSPred consistently outperforms all competing methods across the three datasets. It achieves the highest precision and recall for disease classification, accurately detecting disease cases while minimizing false positives. eSPred also excels in normal classification, particularly in the Lupus and Lung datasets, where competing methods exhibit markedly lower recall. This balanced performance underscores eSPred’s ability to maintain high sensitivity in disease detection without sacrificing specificity for normal samples. Moreover, eSPred demonstrates robustness across datasets with varying degrees of class imbalance: the Lupus dataset shows moderate imbalance (175 disease vs. 99 normal), the Lung dataset exhibits stronger imbalance (250 disease vs. 68 normal), and the COVID-19 dataset presents an inverse imbalance (55 normal vs. 25 disease). Given computational considerations and the larger size of the Lupus and Lung datasets, a single 70%/30% train-test split was used for all datasets.

**Fig. 2:**
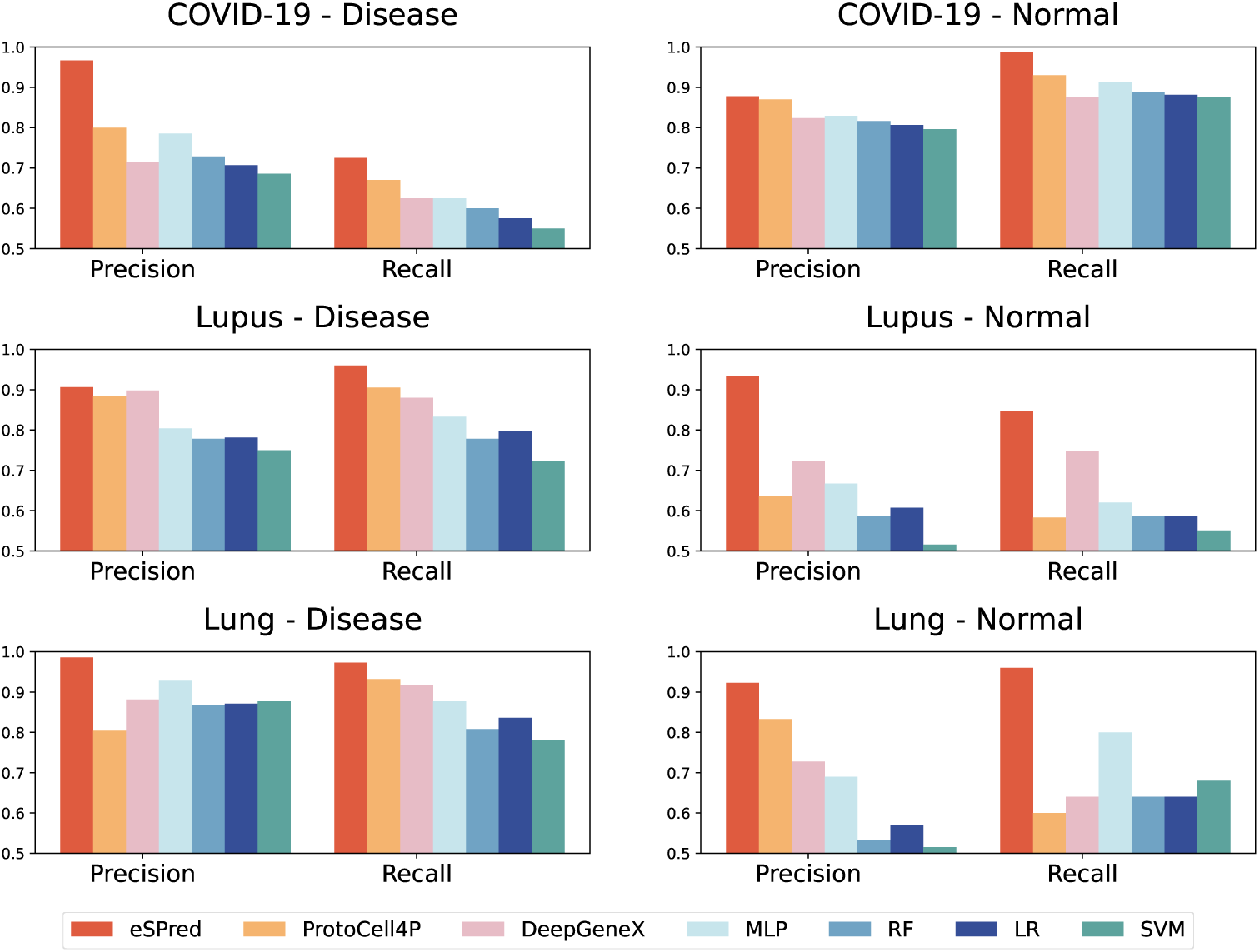
Performance comparison of eSPred and competing methods, measured by precision and recall for disease and normal classification across three datasets: COVID-19, Lupus, and Lung.

**Table 1:** F1 Scores Across Datasets.

| Method | COVID-19 | Lupus | Lung |
| --- | --- | --- | --- |
| scGPT + average cell embedding | 0.772 | 0.637 | 0.784 |
| scGPT + hierarchical classifier | 0.811 | 0.695 | 0.919 |
| scGPT + concatenation + average cell embedding | 0.824 | 0.824 | 0.836 |
| scGPT + concatenation + hierarchical classifier | 0.877 | 0.871 | 0.960 |
| scGPT + pathway + average cell embedding | 0.813 | 0.849 | 0.818 |
| scGPT + pathway + hierarchical classifier | 0.855 | 0.897 | 0.934 |
| **eSPred** | **0.877** | **0.910** | **0.960** |

The superior performance of eSPred can be attributed to several architectural advantages. Unlike traditional approaches that average gene expression across cells or treat cells independently, eSPred preserves cellular heterogeneity through a cell-type-aware grouping strategy, concatenating four cells of the same type to capture shared biological patterns. While ProtoCell4P employs prototype-based learning—which may oversimplify cellular diversity—and DeepGeneX applies feature elimination without biological contextualization, eSPred explicitly integrates gene expression into biologically meaningful pathways. By grounding representations in known functional relationships rather than statistical approximations, eSPred achieves greater classification accuracy and improved clinical relevance.

While these architectural choices intuitively enhance performance, we conducted a systematic ablation study to quantify each individual contribution to eSPred’s predictive accuracy. We progressively incorporated cell-type-aware grouping and pathway integration into the base scGPT model under identical experimental conditions.

Our experimental design systematically evaluated different model variants across three key dimensions: input structure, embedding generation, and subject-level prediction strategy. For input structure, we compared (1) using individual cells as separate inputs (standard in base scGPT and scGPT + pathway variants) and (2) concatenating four cells of the same type into a single input (applied in the scGPT + concatenation variants and eSPred). For cell embedding generation, we compared (1) the [CLS] token embedding, where a special token aggregates information across genes, used in base scGPT and scGPT + concatenation variants; and (2) our pathway-aware embedding method, where information flow is structured according to biological pathways, used in scGPT + pathway and eSPred. Finally, for subject-level prediction, we compared (1) average pooling, where cell embeddings are averaged across all cells within a subject before classification, and (2) hierarchical classifier, where individual cell predictions are aggregated to determine the subject label based on the dominant cell-level classification. This design allowed us to isolate the contributions of input grouping, embedding structure, and prediction strategy to eSPred’s performance improvements.

The base scGPT model with individual-cell input and [CLS] token embedding aggregated through average pooling achieved moderate F1 scores across all datasets (0.772 for COVID-19, 0.637 for Lupus, and 0.784 for Lung). Replacing average pooling with the hierarchical classifier, while maintaining the same [CLS] embedding, consistently improved performance across datasets, confirming the benefit of preserving cellular heterogeneity at the prediction stage.

Next, we evaluated the effect of cell-type-aware concatenation, where four cells of the same type were grouped into a single input during pre-training. Even when combined with [CLS] token embedding and average pooling, concatenation alone led to substantial improvements in F1 scores (0.824 for COVID-19, 0.824 for Lupus, and 0.836 for Lung). Adding the hierarchical classifier on top of concatenation produced further gains, achieving and 0.877, 0.871, and 0.960 across the three datasets, respectively. These results show that adding cell-type information during pre-training, combined with preserving cell-level detail during prediction, leads to substantial performance gains.

We then assessed the impact of pathway-aware embeddings without cell concatenation. The scGPT + pathway variant, using biologically structured embeddings and average pooling, showed notable improvements over the base model, particularly for Lupus (0.849 F1). Using pathway-aware embeddings together with the hierarchical classifier led to better performance across all datasets, showing how biological information can strengthen model representations.

Finally, the full eSPred model, combining cell-type-aware concatenation, pathway-aware embeddings, and hierarchical classification, achieved the best overall performance (0.877 for COVID-19, 0.910 for Lupus, and 0.960 for Lung). These findings highlight three key insights: (1) Hierarchical classification consistently outperforms average pooling, emphasizing the importance of preserving cellular heterogeneity; (2) Incorporating cell-type-aware concatenation during pre-training significantly improves performance, especially in datasets like Lung that have stronger class imbalance; and (3) Pathway integration introduces distinct improvements that enhance the model’s performance in ways not captured by cell-type-aware grouping or hierarchical classification alone.

### eSPred improves cell type representation through cell-type-aware embeddings

To demonstrate the impact of the cell-type-aware grouping strategy in eSPred, we compare three approaches across all datasets: the base scGPT model without grouped inputs, scGPT with grouped inputs, and the full eSPred framework combining grouping with pathway integration. Figure 3 visualizes the resulting cell embeddings using UMAP, highlighting the cellular landscape across the Lupus, Lung, and COVID-19 datasets. These visualizations show a clear improvement in cell type separation when using the grouping (concatenation) strategy incorporated in eSPred, compared to the non-grouped baseline in the base scGPT model.

**Fig. 3:**
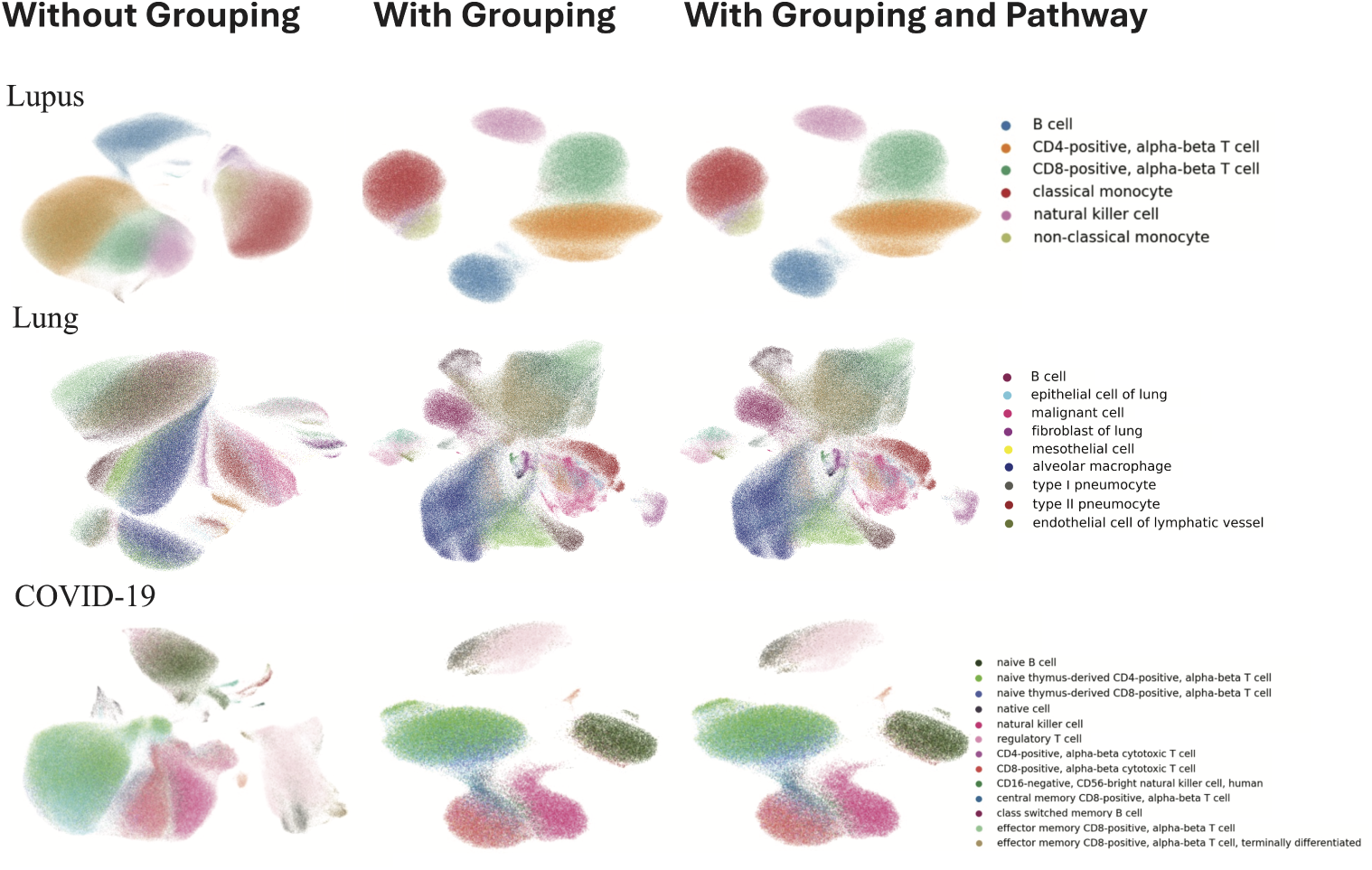
UMAP visualization of cell embeddings across three single-cell RNA sequencing datasets. Each row represents a different dataset (Lupus, Lung, and COVID-19), while columns show three different modeling approaches: base scGPT without grouping (left), scGPT with grouping (middle), and eSPred with both grouping and pathway integration (right). Colors represent different cell types as indicated in the legends.

In the Lupus dataset (top row), embeddings without grouping show extensive overlap among immune cell types, particularly between CD4-positive T cells, CD8-positive T cells, and natural killer cells. Classical and non-classical monocytes also blend without clear separation. Applying cell-type-aware grouping reorganizes these populations into distinct, non-overlapping clusters, with B cells, T cells, and monocytes clearly delineated. Adding pathway integration further stabilizes these boundaries without altering the overall embedding structure. In the Lung dataset (middle row), ungrouped embeddings show substantial mixing among epithelial cells, pneumocytes, malignant cells, and stromal populations. Grouping improves resolution, separating B cells, alveolar macrophages, malignant cells, and various epithelial subtypes into distinct clusters. Pathway integration reinforces these structures, preserving tight and biologically coherent clusters. The COVID-19 dataset (bottom row) shows the most dramatic improvement. Without grouping, T cell subtypes and natural killer cells form a diffuse, heterogeneous cloud. After grouping, naive, regulatory, memory, and effector T cell populations become clearly separated. While pathway integration causes only subtle visual changes, it adds biological coherence to the embedding space, enhancing interpretability and supporting improved downstream prediction.

Overall, these results show that eSPred’s grouping strategy substantially enhances the resolution of cell type embeddings across diverse biological contexts. Although the visual effect of pathway integration is modest, it introduces meaningful biological structure by guiding information flow through known pathways, ultimately contributing to improved classification accuracy and biological interpretability.

### eSPred reveals pathway-level insights in COVID-19

We applied eSPred to the COVID-19 dataset to uncover pathway-level insights into disease mechanisms. While the dataset contained 1,200 HVGs after preprocessing, only 491 genes were mapped to the Reactome pathway database and incorporated into our analysis. These genes were organized into 126 pathways in the first hidden layer and 27 subpathways in the second hidden layer of our neural architecture. As shown in Figure 4, among all pathways, signal transduction emerged as the topranked pathway by importance. This pathway transmission of extracellular signals to intracellular targets, regulating critical immune and stress responses such as cell proliferation, apoptosis, and cycle arrest. During coronavirus infection, SARS-CoV-2 has been shown to manipulate host signaling pathways; for instance, its spike (S) protein upregulates cyclooxygenase-2 (COX-2), a cytokine-regulated enzyme involved in inflammation [70, 71], and activates protein kinase C alpha (PKC*α*), triggering the Raf/MEK/ERK signaling cascade [72–74].

**Fig. 4:**
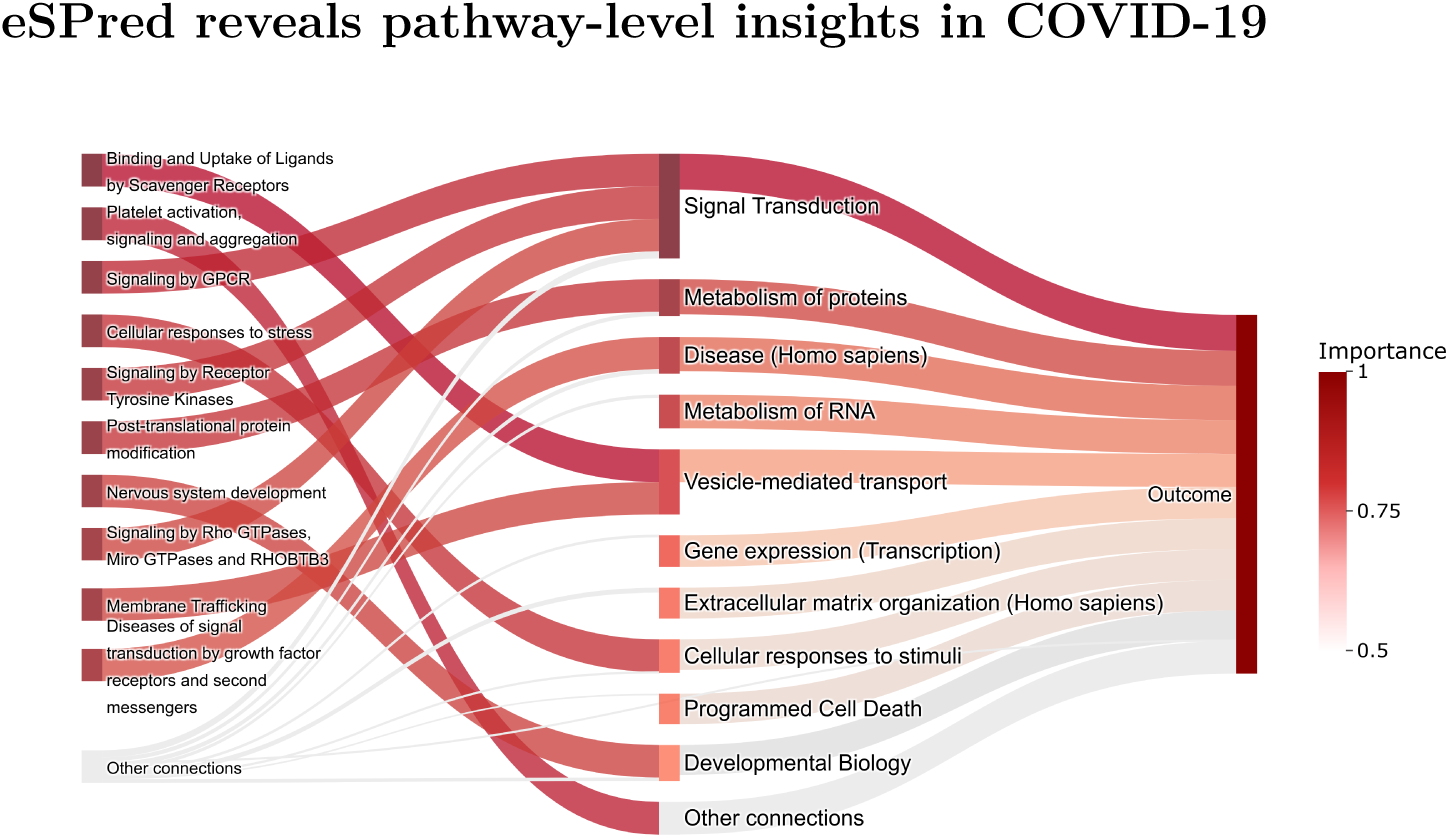
Top ranked pathways identified by eSPred in COVID-19 analysis. The flow chart shows importance scores of key pathways, with immune system, protein metabolism, and signal transduction pathways emerging as critical mechanisms distinguishing COVID-19 patients from healthy individuals. Color intensity indicates the relative importance of each pathway in the classification task.

The immune system pathway ranked second, underscoring its central role in the host response to SARS-CoV-2. Following viral entry, innate and adaptive immune responses are rapidly activated. While early pandemic studies recognized the immune system’s involvement in disease progression, more recent work [75–78] has revealed profound immune dysregulation, shedding light on mechanisms underlying both acute illness and post-acute COVID-19 syndromes.

The third most important pathway, protein metabolism, regulates the synthesis, folding, modification, and degradation of proteins. SARS-CoV-2 infection disrupts these processes by binding the viral spike protein to ACE2 receptors, particularly in lung epithelial cells [79]. Metabolomic studies have further shown that infection alters lipid metabolism and kynurenine pathways, correlating with elevated IL-6 levels and renal dysfunction [80, 81], suggesting that viral replication exploits host lipid metabolism to disrupt protein homeostasis.

These results align with prior experimental findings. Protein–protein interaction studies have confirmed that the SARS-CoV-2 S protein binds ACE2 to enter host cells, impairing enzymatic functions [82, 83]. Changes in lipid metabolism, as observed in metabolomic profiles, offer a biochemical explanation for the model’s emphasis on protein metabolism: hijacking of host lipid pathways supports viral membrane formation and replication, contributing to broader metabolic disruption [81, 84]. Beyond the top three pathways, eSPred identified several additional processes—including cell cycle regulation, transcriptional control, extracellular matrix organization, RNA metabolism, and programmed cell death—as contributing to disease classification. Notably, cytokine signaling within the immune system emerged as a key secondary pathway, consistent with the role of cytokine storms in severe COVID-19, sepsis, HLH, and other hyperinflammatory conditions. Together, these findings highlight eSPred’s ability to uncover biologically coherent, pathway-level mechanisms underlying COVID-19 progression.

#### eSPred highlights the significance of secondary pathways associated with SLE

We applied eSPred to Systemic Lupus Erythematosus (SLE), a chronic autoimmune disease associated with widespread inflammation and multi-organ damage. Out of the initial 1,200 HVGs, 502 were mapped to the Reactome pathway database. These were organized into 128 pathways in the first hidden layer and further grouped into 28 sub-pathways in the second layer. We uncovered a strong predictive association between SLE and disruptions in the O_2_/CO_2_ exchange in erythrocytes pathway. Disruptions in erythrocyte function, particularly involving genes regulating gas exchange, appear significant. In particular, downregulation of the SLC4A1 gene, which encodes the Band 3 protein critical for anion exchange and CO_2_ transport in red blood cells, has been observed in SLE patients [85]. This impairment may contribute to the anemia and fatigue commonly seen in SLE. Additionally, oxidative stress, driven by elevated reactive oxygen species (ROS), can modify erythrocyte components, increasing autoantigen production and promoting autoimmune responses [86].

We also identified strong involvement of G protein-coupled receptor (GPCR) signaling pathways in SLE. GPCRs are critical regulators of immune cell function. In particular, G protein-coupled estrogen receptor 1 (GPER1) activation by estrogen has been shown to enhance monocyte activation in SLE, increasing the expression of pro-inflammatory cytokines such as MCP-1 and TNF-*α* [87]. Furthermore, bioinformatics analyses of B-cell transcriptomes have revealed differential expression of genes involved in GPCR-mediated lysophosphatidic acid (LPA) signaling, pathways known to regulate immune activation and contribute to autoimmune pathology [88].

#### eSPred identifies key pathways associated with lung cancer

In the Lung disease dataset, eSPred analyzed 735 genes mapped to the Reactome pathway database out of the 1,200 HVGs identified during preprocessing. These Reactome-mapped genes were organized into 128 pathways in the first hidden layer and 29 subpathways in the second hidden layer, providing the most extensive pathway coverage among our three datasets. Our eSPred model identified immune system, protein metabolism, and signal transduction pathways as the most influential for classification outcomes, consistent with findings from the COVID-19 dataset. However, our analysis also highlighted additional pathways critical for lung disease stratification.

Among these, the vesicle-mediated transport pathway emerged as a major contributor. Vesicle-mediated transport facilitates the movement of substances within membrane-bound vesicles and plays a vital role in maintaining cellular homeostasis. Prior studies have shown that genes associated with vesicle-mediated transport (VMTRGs) are significant prognostic markers and are closely linked to tumor immunity in lung adenocarcinoma [89, 90]. Dysregulated expression of VMTRGs has been correlated with the activation of key oncogenic signaling pathways, such as Notch and p53, aligning with our findings on secondary pathway importance. The hemostasis pathway also demonstrated a significant role in lung disease classification. Subclinical activation of the hemostatic system is frequently observed in lung cancer patients, characterized by systemic hypercoagulability and regulatory disruption [91]. Hemostasis contributes to tumor progression through various mechanisms, including cancer cell invasion, angiogenesis, immune evasion, and metastasis, making it a critical pathway for disease advancement and a potential therapeutic target [92].

An additional notable finding was the importance of extracellular matrix (ECM)-related pathways. The ECM, composed of proteins such as collagens, fibronectins, laminins, and proteoglycans, provides structural support and regulates key cellular functions like adhesion, migration, and differentiation. In lung adenocarcinoma (LUAD), overexpression of the hyaluronan receptor HMMR has been linked to inflammatory signatures and poor prognosis; attenuation of HMMR reduces tumor initiation and metastasis [93]. Recent studies have further identified ECM-related gene signatures that correlate with LUAD patient survival, immune infiltration, and therapeutic response [94, 95]. Additionally, upregulation of COL10A1 has been associated with ECM remodeling and increased tumor invasiveness in non-small cell lung cancer (NSCLC). By integrating signals from primary (vesicle-mediated transport, hemostasis) and secondary (ECM-related) pathways, eSPred offers a comprehensive insight into the molecular drivers of lung cancer. These findings reinforce known associations while uncovering new aspects of pathway dysregulation relevant to disease classification and management.

### eSPred model achieves fine-grained disease classification

To explore eSPred’s ability to distinguish disease subtypes beyond binary classification, we analyzed subject-level prediction patterns in the lung dataset. Specifically, for each subject, we first obtained cell-level prediction logits—the raw model outputs before softmax transformation—and then averaged these logits across all cells belonging to that subject. The lung dataset includes patients with various lung diseases—chronic obstructive pulmonary disease (COPD), non-small cell lung carcinoma (NSCLC), lung adenocarcinoma (LUAD), and squamous cell lung carcinoma (LUSC)—alongside healthy controls. Although the model was trained only on binary outcomes (disease vs. normal), Figure 7 shows that the analysis of the subject-level logits reveals its ability to differentiate between disease subtypes.

**Fig. 5:**
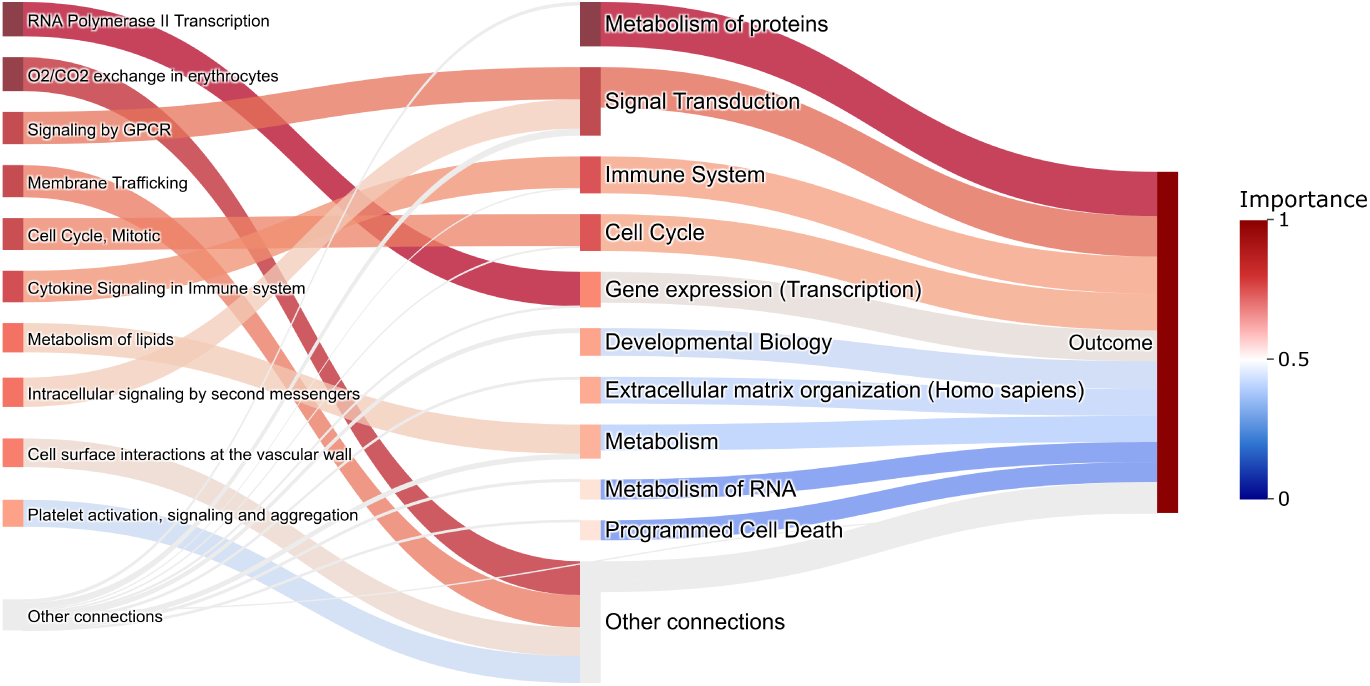
Top ranked pathways identified by eSPred in lupus analysis. The flow chart shows the importance scores of key pathways. Color intensity indicates the relative importance of each pathway in the classification task.

**Fig. 6:**
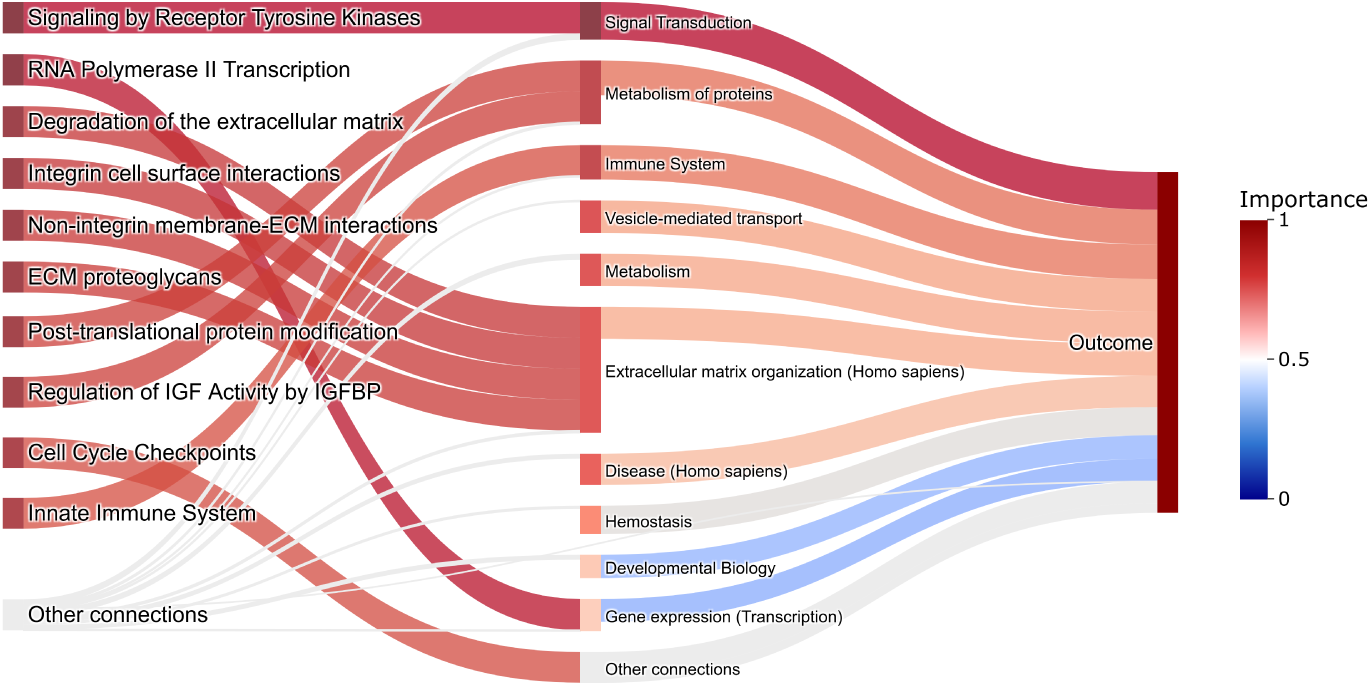
Top ranked pathways identified by eSPred in lung cancer classification. The flow chart shows importance scores of key pathways, including vesicle-mediated transport and hemostasis, alongside immune system and protein metabolism pathways. Color intensity indicates the relative importance of each pathway in the classification task.

**Fig. 7:**
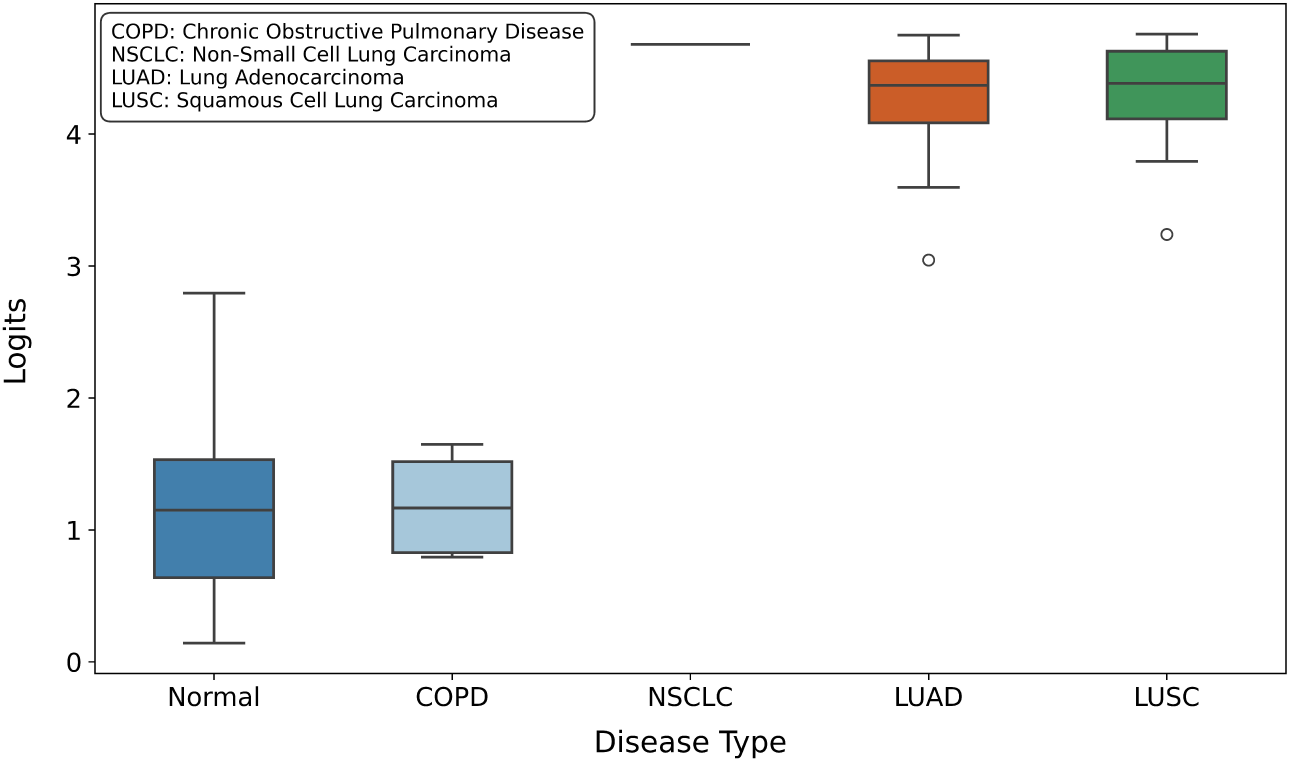
Boxplot showing the distribution of subject-level logit score distributions across four lung disease subtypes and healthy controls.

Carcinoma samples show markedly higher subject-level logits, with NSCLC patients exhibiting the highest average value, followed by LUAD and LUSC. In contrast, normal samples display much lower logits, and COPD cases have the lowest average. While the model clearly distinguishes malignant from non-malignant cases, the overlap between COPD and normal profiles suggests that chronic diseases with milder immune alterations are harder to separate. These results highlight eSPred’s capacity for fine-grained disease classification, even when trained under binary supervision, and underscore its potential for identifying disease subtypes.

## 3 Discussion

We introduce eSPred, a flexible framework for scRNA-seq analysis that enhances foundation model pretraining by incorporating cell-type-specific information and pathway-level biological knowledge. Built on the transformer-based scGPT architecture, eSPred improves predictive accuracy while enabling explicit interpretation of underlying biological processes. eSPred’s design integrates three key innovations. First, a cell-type-aware grouping strategy during pretraining amplifies biologically coherent signals while suppressing technical noise, yielding robust gene–gene and gene–cell representations. Second, a pathway-guided fine-tuning process propagates gene embeddings through a Reactome-informed network structure, aligning the model’s latent space with known signaling pathways without sacrificing predictive performance. Third, a hierarchical two-stage prediction strategy—cell-level classification followed by sample-level aggregation—improving subject-level prediction robustness.

A major strength of eSPred lies in its flexibility and modularity. Although developed using scGPT as the foundation, eSPred can be readily adapted to other large-scale foundation models for single-cell data, such as scBERT, with appropriate data preprocessing and reformatting. Its core innovations—cell-type-aware grouping, pathway-aware fine-tuning, and hierarchical classification—are modular and can be applied to any pre-trained transformer architecture that produces gene-level or celllevel embeddings. This modularity ensures that as new and more powerful foundation models emerge, eSPred’s framework can be easily extended to leverage those advances without requiring fundamental redesign. Our experiments across multiple scRNA-seq datasets demonstrate the transferability and interpretability of eSPred. It achieves accurate predictions while offering pathway-level insights into disease mechanisms and cellular dysfunction.

Despite these advances, pathway-based neural architectures face inherent limitations. First, while leveraging prior biological knowledge improves interpretability, predefined pathways may not fully capture the dynamic and context-specific variability in gene expression across diverse cell types and microenvironments. This mismatch could limit generalization across datasets and biological contexts. Second, although eSPred ranks pathways based on their predictive contribution, it does not perform formal hypothesis testing to assess statistical significance. Unlike traditional enrichment analyses that provide p-values and confidence intervals, eSPred’s pathway rankings may reflect underlying dataset heterogeneity. Additional validation steps, such as permutation testing or external replication, may help further strengthen the biological interpretations.

Several directions could further extend eSPred’s capabilities. While the current cell-type-based grouping strategy is effective, alternative approaches—such as grouping cells by tissue origin or functional state—may offer greater adaptability without requiring full model re-pretraining. However, significant changes to group size would still necessitate retraining and adjustment of embedding dimensions to maintain performance. Beyond binary classification (normal vs. disease), eSPred’s architecture can naturally support multi-class predictions. Replacing the output layer with a multi-class softmax would allow modeling of multiple disease categories or subtypes, particularly when no clear severity hierarchy exists. Coupled with pathway-aware embeddings, this extension could provide a finer resolution of disease heterogeneity. Although batch effects were minimal in our datasets, future versions of eSPred could integrate scGPT’s batch-correction objectives to improve robustness across more heterogeneous scRNA-seq experiments. Enhancing batch alignment would further preserve biological signals while reducing technical variability.

In summary, eSPred demonstrates the potential of foundation model–based approaches for interpretable, scalable, and clinically meaningful single-cell analysis. Its flexible, modular design allows seamless adaptation to future foundation models, paving the way toward robust, personalized predictions across increasingly complex biological datasets.

## 4 Methods

We develop eSPred, an enhanced framework built on the scGPT foundation model, to improve clinical prediction and interpretability in single-cell RNA sequencing data. Our approach extends the original scGPT architecture through three key innovations: (1) incorporating cell-type information during pre-training, (2) introducing a pathway-guided embedding mechanism during fine-tuning, and (3) implementing a hierarchical prediction strategy for subject-level outcomes. These enhancements enable eSPred to generate biologically meaningful representations while maintaining strong predictive performance across diverse disease contexts.

### Enhanced scGPT with cell-type information integration

The original scGPT foundation model applies a transformer architecture to scRNAseq data, treating genes as “tokens” similar to words in language models. For each cell, scGPT takes three main components as input: (1) gene token vectors that assign unique identifiers to each gene, (2) gene expression value vectors representing measured expression levels, and (3) condition token vectors encoding metadata such as treatment status, batch, tissue type, or disease state. These inputs are combined through embedding layers and then processed by a series of transformer blocks, each containing multi-head attention mechanisms and feed-forward neural networks. This architecture allows the model to learn complex dependencies between genes by enabling each gene to attend to all others within the same cell. Using this architecture, scGPT produces two types of embeddings: gene embeddings, which are 512-dimensional vectors capturing gene-level interactions and functional relationships, and cell embeddings, typically derived from a special [CLS] token, which summarize the overall cellular state. These embeddings support a range of downstream applications, such as cell type annotation and perturbation response prediction.

In our work, we enhance the scGPT framework by incorporating cell-type information through a grouping strategy. Rather than processing individual cells independently, we identify cells of the same type—either manually or via clustering—and organize them into groups of four. The gene expression profiles from these four cells are concatenated to form an aggregated representation, providing a more robust input for the model. Let K denote the set of cell-type indices, and let **c** represent a cell in the dataset. For each cell type *k* ∈ K, we define the set of corresponding cells as

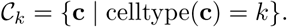

From each set C*_k_*, we form groups of four cells, indexed by *j*, as follows:

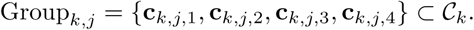

During fine-tuning, we impose an additional constraint: all cells within a group must come from the same subject. This ensures the model captures both cell-type-specific and subject-specific signals.

This cell-type-aware grouping offers two key benefits. First, it improves computational efficiency by processing four cells at once while respecting the scGPT token limit (∼5,000 tokens). Since each cell includes 1,200 HVGs, grouping four cells results in 4, 800 tokens—fully utilizing the model’s input capacity in a single pass. Second, and more importantly, this strategy strengthens biological signal detection. By aggregating expression profiles within the same cell type, we amplify shared biological patterns while mitigating technical noise—such as dropout events, batch effects, and variable capture efficiency—that tends to occur randomly across cells. This averaging effect improves the signal-to-noise ratio, enhancing the model’s ability to distinguish disease-specific patterns during downstream classification tasks.

### Pathway-aware embedding generation

After pre-training with cell-type-aware groups, we introduce a pathway-guided mechanism to generate biologically interpretable embeddings. When a cell’s gene expression profile is processed through the enhanced scGPT model, it produces a gene embedding matrix of size 512 × 1200, where 1200 corresponds to the number of HVGs and each gene is represented by a 512-dimensional vector that captures its functional and contextual properties. Let *E* ∈ ℝ*^G^*^×^*^Q^* represent the gene embeddings matrix from our enhanced pre-trained scGPT model, where *G* is the number of genes (1200) and *Q* = 512 is the embedding dimension. To incorporate biological prior knowledge, we use gene–pathway mappings from the Reactome database to build a hierarchical neural network with *l* levels (typically *l* = 4). Let P*_i_* be the set of pathways at level *i*, and let *M_i_* ∈ {0, 1}^|P^*^i^*^|×|P^*^i−^*^1 |^ be the binary mask matrix encoding the connections between pathways at levels *i* and *i* − 1. The activation of each pathway node *p* at level *i* is computed as:

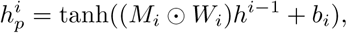

where *W_i_* is a learnable weight matrix, *b_i_* is the bias term, ⊙ denotes element-wise multiplication with the binary mask, and *h*^0^ = *E* is the input gene embedding matrix. The activation function tanh(·) follows the setting used in CellTICS. Each node in the hierarchy acts as an aggregator, combining signals from its associated genes or lower-level pathways. To predict the disease state of a subject, we apply a softmax layer to the top-level pathway activations:

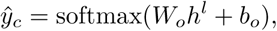

where *h^l^*is the final-level pathway representation, and *W_o_* and *b_o_* are learnable output parameters.

To assess the contribution of each pathway to distinguishing between disease states, we compute pathway-specific importance scores based on differences in pathway activation patterns. Our primary focus is on binary classification tasks (e.g., normal vs. disease). For a given pathway *p* and disease state *s*, we first compute the mean activation vector:

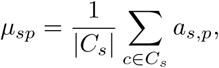

where *C_s_*is the set of cells in state *s*, and *a_c,p_* is the activation vector of pathway *p* for cell *c*. The activation vector *a_c,p_*is the *Q*-dimensional output (where *Q* = 512) produced at pathway node *p* when cell *c* is processed through our hierarchical neural network, i.e., the vector representation obtained after applying the tanh activation function at the corresponding pathway node. These vectors are then normalized using L2 normalization

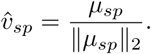

For binary comparisons between disease states *s*_1_ and *s*_2_, the importance score for pathway *p* is defined as the L2 distance between their normalized activation vectors:

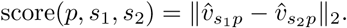

For multi-class classification (e.g., more than two disease states), we follow the approach from CellTICS. For each pathway *p*, we compute the average activation across all disease states and sort them in descending order: *A*_(1)_*_,p_, A*_(2)_*_,p_, …, A*_(_*_S_*_)_*_,p_*, where *S* is the number of states. The pathway importance score is then defined as the difference between the top two average activations:

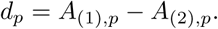

Since a pathway may be present in multiple prediction layers, the final importance score *D_p_* is defined as the maximum *d_p_* value across all paths. Pathways with higher scores exhibit more distinct activation profiles across disease states, indicating their potential as key discriminative features. This approach captures both directional differences and relative activation strengths, enabling the identification of pathways most strongly associated with disease-specific immune states. The resulting scores provide interpretable insights into underlying biological mechanisms.

Our pathway-aware approach offers two key innovations compared to standard foundation models:

#### Biologically-constrained information flow

In the original scGPT architecture, following the transformer design used in models like BERT, a special [CLS] token is prepended to the input sequence for each cell. This token attends to all gene positions during self-attention, functioning as a global aggregator of information. While effective for producing cell embeddings, the [CLS] token operates as a black box—its attention weights are not biologically guided, and it can mix gene information without structural constraints. As a result, it becomes difficult to interpret how specific biological processes contribute to the final embedding or prediction. To address this, our approach replaces the [CLS] mechanism with a biologically grounded alternative: a pathway-guided architecture informed by Reactome. Gene embeddings are propagated through a hierarchical network that mirrors known biological pathways, ensuring that downstream representations are built from biologically meaningful groupings. This structure not only improves interpretability—by enabling direct attribution of predictive signals to specific pathways—but also enforces biologically plausible information flow.

#### High-dimensional pathway representation

Conventional pathway analysis tools, such as CellTICS, typically rely on scalar gene expression values as input. Each gene contributes a single numerical value, limiting the granularity of information passed to the pathway level. In contrast, our framework leverages the full 512-dimensional gene embeddings generated by the foundation model. These rich representations are propagated through the pathway hierarchy, allowing each pathway node to act as a high-capacity aggregator of complex input signals. This enables a more expressive modeling of pathway activity, preserving the nuanced gene-gene and gene-pathway relationships learned during pretraining. As a result, pathway activity scores derived from our model capture subtler biological signals than traditional scalar-based methods, without sacrificing interpretability.

### Hierarchical cell-to-subject outcome prediction

To derive subject-level predictions from cell-level data, we implement a hierarchical prediction strategy. Let N be the set of all subjects, and for each subject *n* ∈ N, let C*_n_* denote the set of cells from that subject. Each cell ***c*** ∈ C*_s_* is processed through our enhanced scGPT model to obtain gene embeddings, which are then passed through a pathway-aware neural network to generate a cell-level prediction *y*^*_c_* ∈ {0, 1}. To aggregate these into a subject-level prediction, we apply a simple majority voting rule:

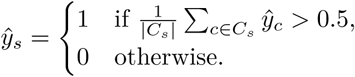

This approach reflects the biological expectation that, while gene expression may vary across cells, the overall immune profile should be consistent with the subject’s clinical state. Aggregating across cells enhances robustness, mitigating the effects of technical noise and outlier cells, and supports reliable subject-level predictions.

### Baseline models

To ensure fair comparisons across all experiments, we used the same training and testing datasets for each model. Specifically, we selected 1,200 highly variable genes (HVGs), and their corresponding expression values served as the standardized input for all baseline models.

#### Logistic Regression

Logistic regression (LR) estimates the probability that a sample belongs to a given class by applying a linear transformation to the input features followed by a sigmoid activation function. For each sample, the input features are defined as the mean gene expression values averaged across all cells. While LR is simple and interpretable, its linear decision boundary limits its ability to capture the complex, non-linear relationships often present in single-cell data. In our experiments, we tuned hyperparameters by varying the regularization penalty ∈ {l1, l2, none}, the solver ∈ {liblinear, lbfgs, saga}, and the maximum number of iterations ∈ {1000, 5000, 10000}.

#### Random Forest

Random Forest (RF) is an ensemble learning method that builds multiple decision trees, each trained on random subsets of the data and features, and aggregates their predictions through majority voting. In our analysis, the input features are the mean gene expression values across all cells in a sample, and the RF model outputs the most frequently predicted class label among the trees. We explored various hyperparameter configurations, including the number of trees ∈ {100, 200, 500}, the maximum depth of each tree ∈ {None, 10, 20}, and the minimum number of samples required to split an internal node ∈ {2, 5, 10}.

#### Support Vector Machine

Support Vector Machine (SVM) is a classification algorithm that constructs an optimal decision boundary by maximizing the margin between classes in the feature space. It supports both linear and non-linear classification through the use of kernel functions. In our experiments, we evaluated both linear and radial basis function (RBF) kernels. Hyperparameter tuning was performed by varying the regularization parameter *C* ∈ {0.01, 0.1, 1, 10}, the tolerance for convergence tol ∈ {1e-4, 1e-3, 1e-2}, and the maximum number of iterations max iter ∈ {1000, 5000, 10000}.

#### Multi-Layer Perceptron

Multi-Layer Perceptron (MLP) is a feedforward neural network that models complex, non-linear relationships using multiple fully connected hidden layers. In our setup, the model receives individual cell-level gene expression profiles as input and outputs celllevel predictions. These predictions are then aggregated via majority voting to derive subject-level classifications. We experimented with network architectures containing 1 to 3 hidden layers, with the number of neurons per layer set to {64, 128, 256}. We further tuned the learning rate ∈ {1e-4, 1e-3, 1e-2} and dropout rate ∈ {0.2, 0.5} to improve model generalization and convergence.

#### ProtoCell4P

ProtoCell4P [55] is a prototype-based neural network designed for patient-level classification using scRNA-seq data. The model takes individual cell expression profiles as input and encodes them into a low-dimensional embedding space, where it learns a set of representative cell prototypes. Each cell is evaluated based on its similarity to these prototypes, and corresponding relevance scores are computed to determine its contribution to the subject-level prediction. Final predictions are obtained by aggregating cell-level signals across all cells from a given subject. While ProtoCell4P leverages cell type information to enhance interpretability, its prototype-based architecture may oversimplify complex cellular interactions, potentially limiting its flexibility in heterogeneous clinical datasets. For our implementation, we used the Python code available at https://github.com/Teddy-XiongGZ/ProtoCell4P. We performed hyperparameter tuning by varying the hidden dimension size h dim ∈ {64, 128, 256}, learning rate lr ∈ {1e-4, 5e-4, 1e-3}, and the number of prototypes n proto ∈ {10, 20, 50}.

#### DeepGeneX

DeepGeneX [53] is a framework for gene feature selection and patient classification based on scRNA-seq data. It constructs subject-level representations by averaging gene expression values across all cells within a sample. To identify informative genes, DeepGeneX employs a feature elimination strategy: one gene is shuffled at a time, and the resulting increase in binary cross-entropy loss is used to assess that gene’s importance. This iterative process ranks genes by their contribution to prediction accuracy and progressively eliminates those with lower importance scores. While DeepGeneX effectively reduces input dimensionality and highlights key predictive genes, its focus on marginal gene importance may limit its ability to capture complex gene–gene interactions or higher-order dependencies—factors that could be critical for accurate clinical outcome prediction. We downloaded Python codes for DeepGeneX from their Github: https://github.com/gujrallab/biomarkers. In our experiment, we utilized the hyperparameter tuning methods provided in their code to optimize the model. This tuning and feature elimination process reduced the gene set from 1,200 to 60, with the resulting deep neural network model consisting of 3 layers and approximately 16,000 parameters.

### Datasets

#### Data preprocessing

The scRNA-seq data were preprocessed using the Scanpy library. First, genes expressed in fewer than 10 cells were removed to eliminate low-information features and reduce noise. Next, total counts per cell were normalized to a target sum of 10,000 to account for differences in sequencing depth across cells. The normalized expression values were then log-transformed using the natural logarithm (*log*(1 + *x*)) to stabilize variance and mitigate the influence of highly expressed genes. Finally, the top 1,200 HVGs were selected based on their dispersion, focusing the analysis on features that capture the most biologically meaningful variation across cells.

#### COVID-19 dataset

This dataset comprised 80 subjects, including 25 COVID-19 patients and 55 normal controls. A total of 422,220 single cells were profiled across 1,200 genes, spanning 30 annotated cell types. The dataset was split into training and testing sets using a 70/30 ratio. The training set included 56 subjects (17 COVID-19 patients and 39 controls), while the testing set included 24 subjects (8 COVID-19 patients and 16 controls). As part of preprocessing, cells that did not belong to groups with at least four cells per subject were excluded. Notably, granulocytes were entirely removed from the analysis, as they did not meet this minimum threshold required for our cell clustering approach. As a result, the number of cell types was reduced from 30 to 29. The cell type distribution in both the training and testing sets maintained a similar pattern to the original dataset.

#### Lupus dataset

This dataset included 274 subjects: 175 with systemic lupus erythematosus (SLE) and 99 normal controls [96]. It comprised 1,263,676 single cells profiled across 10,000 genes, annotated into 11 immune cell types. We selected the top 1,200 HVGs during preprocessing and split the data into training and testing sets using a 70/30 ratio. The training set included 191 subjects (125 SLE and 66 controls), and the testing set contained 83 subjects (50 SLE and 33 controls). The final dataset included 1,262,878 cells—887,052 in training and 375,826 in testing. The slight reduction in cell count was due to our cell grouping strategy, which retained only cells that could be clustered into sets of four within the same cell type. All 11 cell types were preserved, with their proportions consistent across both sets.

#### Lung dataset

This dataset included 318 subjects, consisting of 250 disease cases and 68 normal controls [97]. Among the disease cases were 172 with lung adenocarcinoma (LUAD), 45 with squamous cell carcinoma (LUSC), 15 with non-small cell lung carcinoma (NSCLC), and 18 with chronic obstructive pulmonary disease (COPD). A total of 1,283,972 single cells were profiled across 17,797 genes and annotated into 33 immune cell types. We selected 1,200 highly variable genes to reduce dimensionality while retaining meaningful biological variation. The data was split into training and testing sets using a 70/30 ratio, resulting in 220 training subjects (177 disease, 43 control) and 98 testing subjects (73 disease, 25 control). Disease subtypes were proportionally distributed across both sets. After preprocessing, the total cell count was slightly reduced to 1,281,410. This was due to our requirement that cells be grouped in sets of four by type—cells that didn’t fit into such groups were excluded. All 33 cell types were preserved, and their distributions remained consistent between the training and testing sets.

We evaluate model performance using precision, recall, and F1-score, which are standard metrics for binary classification tasks. Precision reflects how often positive predictions are correct, indicating the model’s ability to avoid false positives. Recall measures how well the model identifies all true positive cases, highlighting its sensitivity to relevant signals. The F1-score combines precision and recall into a single value, offering a balanced view of overall performance. These metrics are calculated by comparing predicted class labels to the ground truth labels in the test set.

## 5 Availability of data and materials

The COVID-19 and lupus datasets are publicly available and can be accessed via https://figshare.com/articles/dataset/pbmc_raw_h5ad_gz/21688029. The lung dataset is available through the CellxGene portal: https://cellxgene.cziscience.com/collections/edb893ee-4066-4128-9aec-5eb2b03f8287. An open-source implementation of eSPred is available on GitHub at https://github.com/llin-lab/eSPred.

